# Patterns of DNA variation between the autosomes, the X chromosome and the Y chromosome in *Bos taurus* genome

**DOI:** 10.1101/2020.01.21.901348

**Authors:** Bartosz Czech, Bernt Guldbrandtsen, Joanna Szyda

## Abstract

The new ARS-UCD1.2 assembly of the bovine genome has considerable improvements. That might be assumed that a more accurate identification of patterns of genetic variation can be achieved with it. We explored differences in genetic variation between autosomes, the X chromosome, and the Y chromosome. In particular, densities of variants, annotation, lengths (only for InDels), nucleotide divergence, and Tajima’s D statistic between chromosomes. Whole-genome DNA sequences of 217 individuals representing different cattle breeds were examined. The analysis included the alignment to the new reference genome and variant calling. 23,655,295 SNPs and 3,758,781 InDels were detected. In contrast to autosomes, both sex chromosomes had negative values of Tajima’s D and lower nucleotide divergence. That implies a correlation between nucleotide diversity and recombination rate, which is obviously reduced for sex chromosomes. Moreover, accumulation of nonsynonymous mutations on the Y chromosome could be associated with loss of recombination. Also, the relatively lower effective population size for sex chromosomes leads to a lower expected density of variants.

## Introduction

DNA variation refers to differences in DNA sequence among individuals. Decreasing costs and reducing time of whole-genome sequencing using Next-Generation Sequencing (NGS) technology bring the opportunity to sequence many samples. The analysis of differences in DNA variation between autosomes and sex chromosomes plays an important role in understanding the evolution of chromosomes. In terms of studying variation, cattle is an interesting model, since for a number of generations it has undergone strong artificial selection toward increased levels of production. In addition, modern cattle is composed of breeds with distinct phenotypic characteristics. We used the new ARS-UCD 1.0.25 assembly ^1^. However, this assembly, like the UMD3.1^2^, does not contain the bovine Y chromosome (BTY), so most analyzes ignored the Y chromosome as a consequence it has been omitted from most studies, so that only a few associations between BTY and phenotypes were identified in humans and only one in livestock (pigs, source AnimalQTdb www.animalgenome.org/cgi-bin/QTLdb/index). The 1000 Bull Genomes Project ^3^ added the Y chromosome assembly from Btau 5.0.1 to the ARS-UCD, creating the ARS-UCD1.2_Btau5.0.1Y assembly. Thereby it becomes feasible to work with all nuclear bovine chromosomes.

According to the ARS-UCD1.2_Btau5.0.1Y reference genome assembly, the bovine genome spans 2,759,153,975 bp with 30,278 genes (GeneBank assembly accession: GCA_002263795.2 and GCA_000003205.6) and consists of 29 autosomes and two sex chromosomes – X, and Y. The hemizygous Y chromosome in *Bos taurus* is short and contains only few genes. In contrast, the bovine X chromosome (BTX) contains many more genes and its length is similar to the autosomes. Chromosome 1 is the longest bovine chromosome, spanning 158,534,110 bp with 1,218 genes, while chromosome 25 is the shortest bovine chromosome (not much shorter than BTY), spanning 42,350,435 bp with 1,611 genes. BTY contains the lowest number of genes (206), while chromosome 23 contains the highest number of genes (1,708). Clearly, chromosome length is not linearly related to the number of genes. Moreover, 29 autosomes are acrocentric, while both sex chromosomes (X and Y) are submetacentric. Since recombination that generates new combinations of alleles is characteristic to autosomes, on sex chromosomes, this phenomenon is reduced in males to only small homologous regions shared between BTX and -Y, called the pseudoautosomal regions (PARs). BTY is poorly characterized because it is difficult to sequence due to the occurrence of a high proportion of repetitive sequences ^4^. There is a general tendency toward degeneration of functional elements.

Differences in genetic variation patterns arise from variable recombination rates and mutation rates, genetic drift, demography, selection and population history. Therefore the focus of our study was on the comparison of patterns of genetic variation between autosomes, the X chromosome and the Y chromosome in the context of bovine genome.

## Results

### Alignment to the reference genome

The quality of alignment was expressed by the total percent of mapped reads and the percent of properly paired mapped reads (both reads are mapped close to each other in opposite directions on the same chromosome). The percent of mapped reads for each individual was very high and ranged from 91.4% to 99.9% with mean 99.7% (±0.6) and very similar mode 99.9%. The percent of properly paired mapped reads was also high and varied between 89.3% and 99.1% with mean 97.1% (±1.7) and mode 97.9%.

Average genome coverage was calculated separately for each individual and ranged from 5.28 to 46.79 with mean and mode both equal to 25 (Figure 1). Individuals average genome coverage less than 15, while two individuals have average genome coverage above than 40. The individual with the lowest average genome coverage (5.28) contains 99.6% of mapped reads and 97.8% of properly paired mapped reads. The individual with the highest average coverage (46.79) contains 99.9% of mapped reads and 97.9% of properly paired mapped reads.

**Figure 1.**
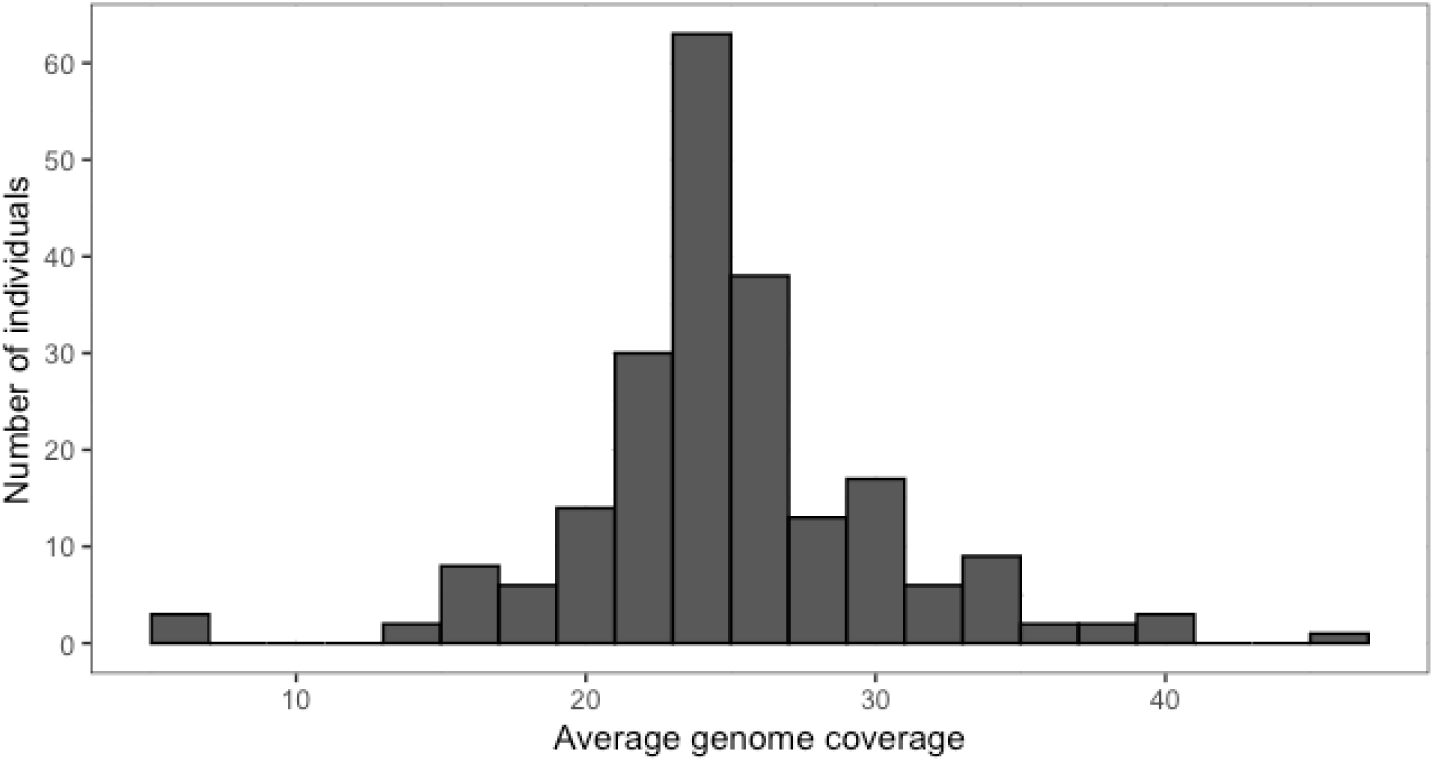
Average genome coverage.

### Variation

Over all individuals 27,414,076 variants were identified. Of these, 86.3% were SNPs, and 13.7% were InDels. BTA1 contained the highest number of variants (1,689,556), while the BTY contained the lowest number of variants (49,591) (Figure 2). The total number of SNPs was 23,655,295, 0.9% of the total genome length. Figure 2 presents the distribution of SNPs. BTA1 contained the highest number of SNPs (1,455,295), but the highest SNP density, expressed by the proportion of the number of SNPs to length, was highest for BTA25 (2.6%). BTY was characterized by the lowest number of SNPs (41,500) and the lowest SNP density (0.1%). 3,758,781 of InDels were identified. BTA1 had the highest number of InDels (234,261), whereas BTY had the lowest number of InDels (8,091) (Figure 2).

**Figure 2.**
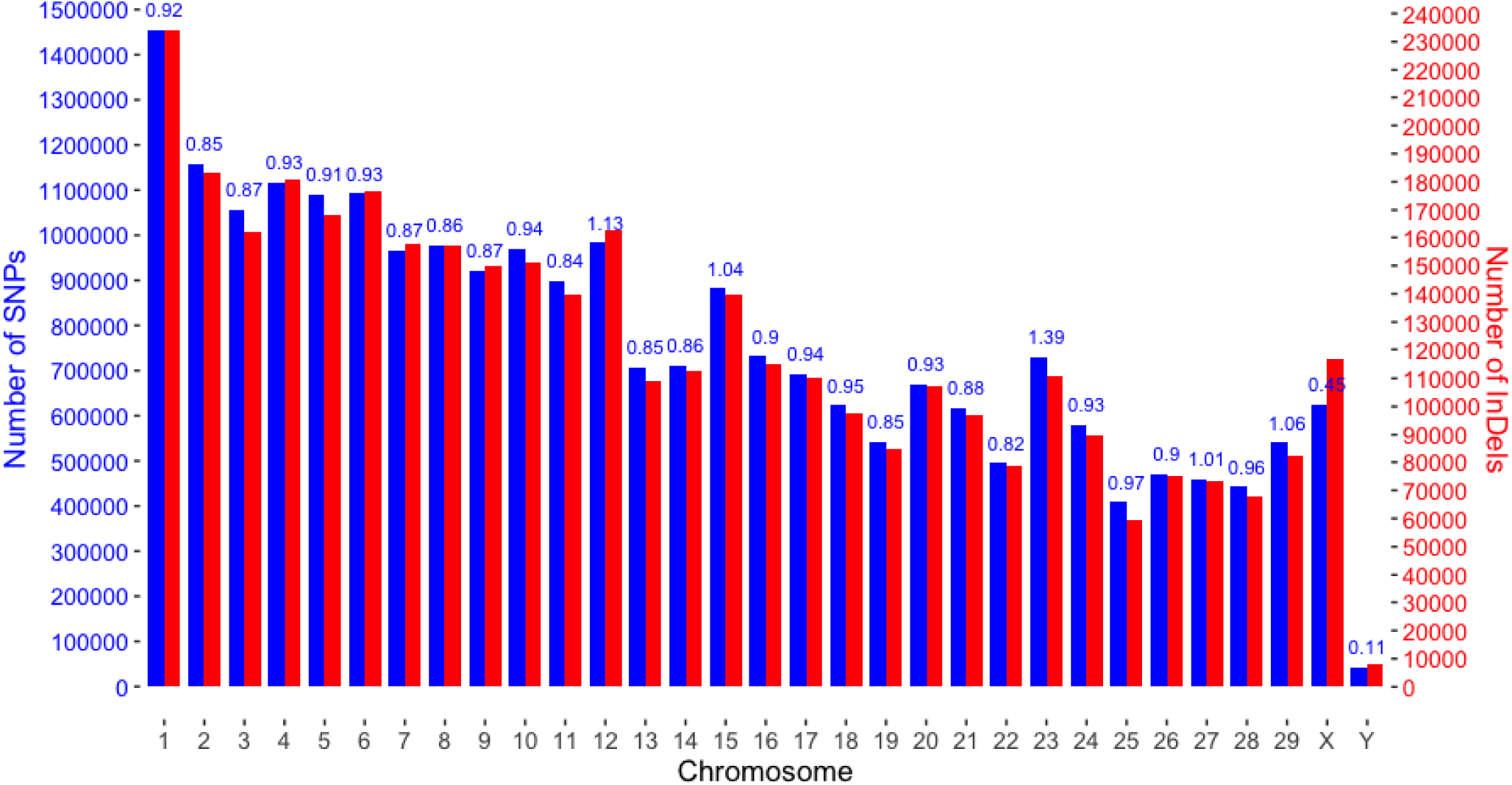
The distribution of variants.

The length of InDels varied between 1 bp and 281 bp on autosomes, from 1 bp to 233 bp on BTAX, and from 1 bp to 156 bp on BTY. The most frequently observed length of InDels was one bp, but the median length was two bp (Figure 3). However, InDel median length differed significantly (*P* < 0.001) among autosomes vs. BTX and autosomes vs. BTY. Neither autosomes’ nor sex chromosomes’ InDel length distributions do not follow the uniform distribution (*P* < 0.001). The percent of InDels at least 10 bp long was 12%, 19%, and 98% for autosomes, BTX and BTY, respectively.

**Figure 3.**
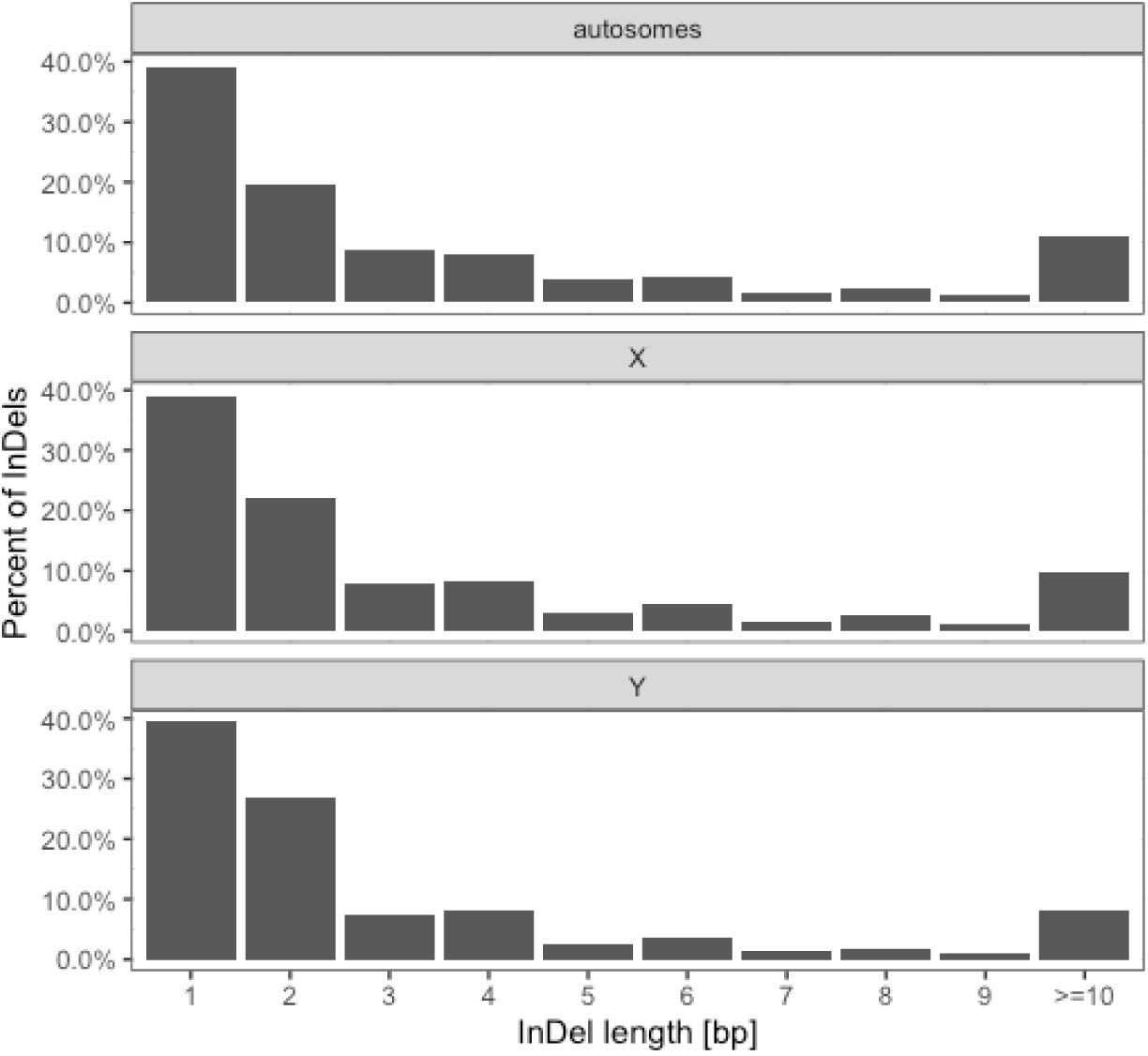
Length distribution of deletions and insertions for autosomes and sex chromosomes.

### Distribution of Polymorphisms

The analysis of variant distribution was done by counting the number of variants within 100 kbp non-overlapping windows (Figure 4). The number of SNPs per 100 kbp bin was heterogeneous across windows within the same chromosome (*P* < 0.001). The largest number of SNPs in one window (16,184) was observed on BTA12. On the other hand, on four chromosomes we identified windows without SNPs: BTA8 (1 window), BTA9 (8), BTA10 (8), and BTY (58). All of the windows were annotated as intergenic. However, 27 out of the total 58 windows on BTY corresponded to gaps in the reference genome marked by stretches of Ns. Conversely, windows with the number of SNPs exceeding 10,000 were found on BTA4 (one window with 10,838 SNPs) 8% of window length overlapped with genes, BTA12 (four windows with 10,431, 11,487, 15,097 and 16,186 SNPs) in 3% of windows length overlapped with genes, and BTA23 (three windows with 11,706, 12,452 and 13,229 SNPs) in which 100% of windows length overlapped genic regions. Interestingly, the functional annotation showed that four out of the eight SNP-rich windows overlapped with lncRNA genes.

**Figure 4.**
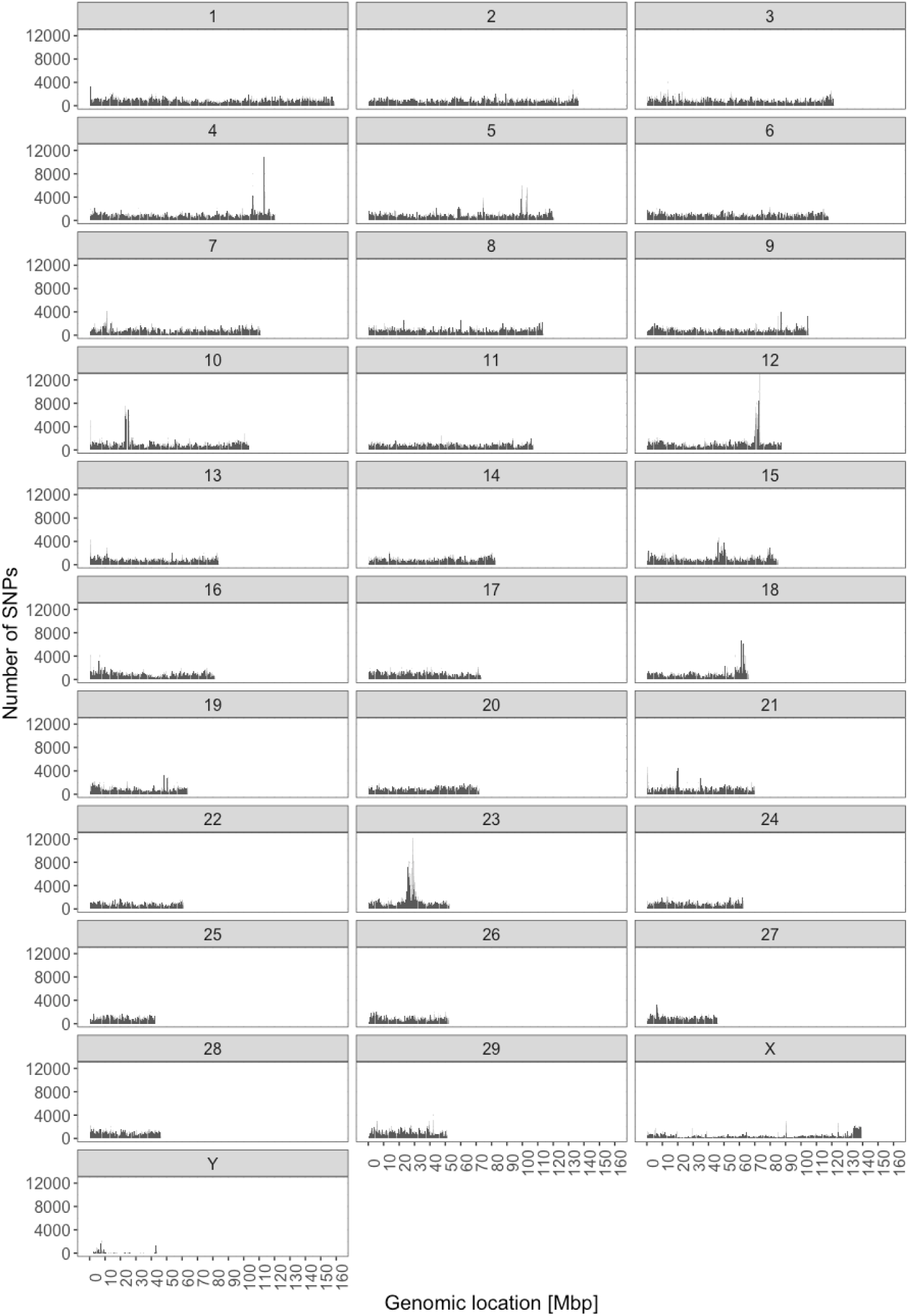
The distribution of SNPs across chromosomes.

The highest overall number of InDels was found on BTA23 (2,930 InDels). InDel density (Figure 5) was not uniform across the genome (*P* < 0.001). We identified 122 windows without InDels – 7 on BTA9, 8 on BTA10, and 107 on BTY. All the windows on BTA7 and BTA8 were intergenic. On BTY, 64 of the windows were intergenic, whereas 35 windows corresponded to gaps in the reference assembly. However, 8 windows intersected 10 genes. 1% to 13% of these windows overlapped with genic sequence. Windows harbouring more than 2,000 InDels were found on BTA4 (one window with 2,162 InDels), BTA12 (two windows with 2,405 and 2,792 InDels), and BTA23 (three windows with 2,022, 2,258, and 2,930 InDels). Each of these windows overlapped with genes comprising from 33% of a window length to 100% of length, both on BTA23. As for SNP-rich windows, three out of the six InDel-rich windows were overlapped with lncRNA genes.

**Figure 5.**
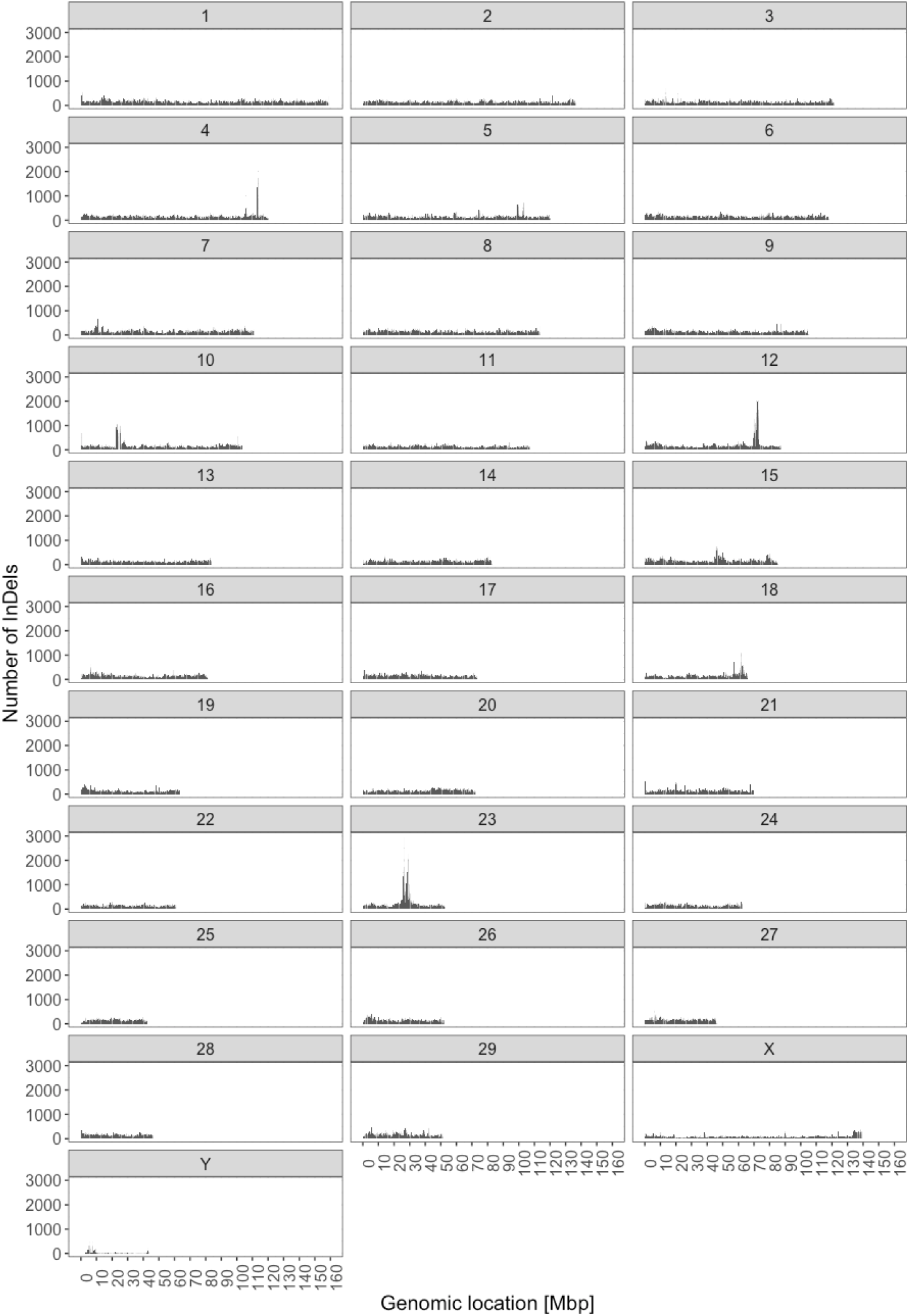
The distribution of InDels across chromosomes.

### Variant annotation

The annotation of variants expressed as the percent of polymorphic bases located in intergenic or genic regions in relation to their length was summarized on Figure 6. Regardless of annotation, autosomes had the highest and BTY had the lowest density of polymorphic sites. On each autosome, intergenic variants were the most common. Between 1.03% and 1.25% of intergenic bases were annotated as polymorphic. In genes, between 0.86% and 1.17% of bases were polymorphic. On BTX, the 0.53% of intergenic and 0.54% of genic were polymorphic. In contast, on BTY 0.2% of genic and 0.1% intergenic bases were polymorphic.

**Figure 6.**
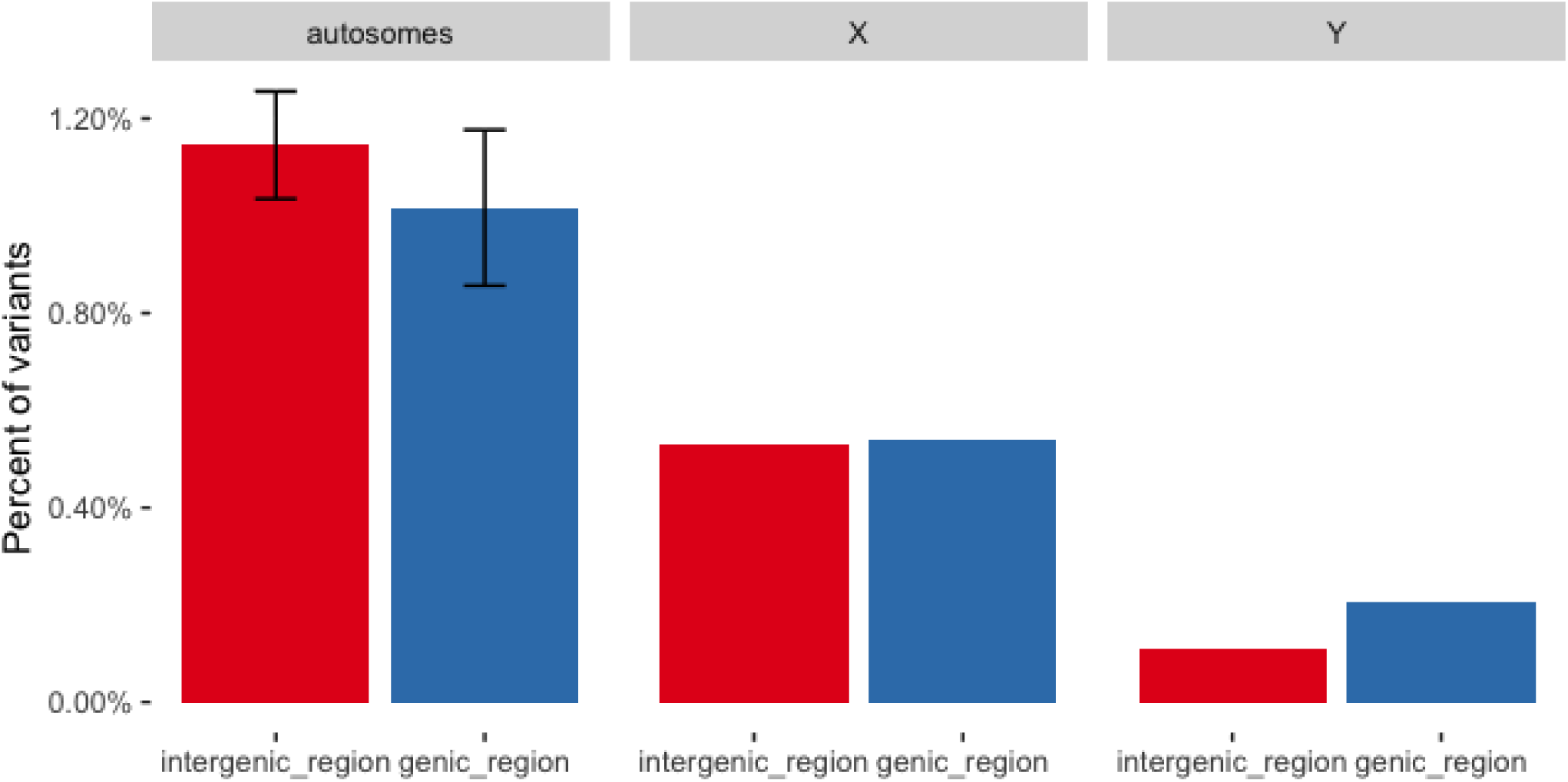
The annotation of variants to genic and intergenic regions.

Focusing only on genes, on autosomes and BTX we observed the highest percent of variants was located in introns (41.8% ± 2.9% and 42.6%, respectively) and in non-coding transcripts (41.7% ± 2.4% and 41.7%, respectively). The Y chromosome showed the highest percent of genic variants annotated 5000 bp upstream of the 3’ end of gene start (25.3%) and 5,000 bp downstream of the 5’ end of gene end (24.8%). It was accompanied by a proportion of intronic and non-coding transcript variants being by 20% lower than in BTA and BTX. Moreover, variants with high (splice acceptor, splice donor, stop gained, frameshift variant, stop lost, start lost) and moderate (missense) impacts on proteins occurred more frequently on BTY than on BTA and BTX, while the latter had the lowest percent of variants with high or moderate impacts. Interestingly, on BTY 0.11% of genic variants are frameshift, which is much higher than on BTA (0.02%) and BTX (0.01%). The same tendency was observed for stop gained variants, which constitute 0.03% of BTY, but only 0.004% of BTA and 0.003% of BTX.

### Population genetics

The ratio of non-synonymous to synonymous SNPs (*K*_*a*_*/K*_*s*_ ratio) was the highest for BTY (2.00) and the lowest for BTX (0.62). Autosome averaged ratio was also below unity (0.79 ± 0.15) and varied between 0.57 for BTA22 and 1.10 for BTA4, BTA15, BTA18, and BTA23. Tajima’s *D* statistic significantly differed between all three chromosome groups (*P* < 0.001) and was positive for autosomes and negative for both sex chromosomes (Figure 7). Nucleotide diversity had the highest median for autosomes, followed by BTX and BTY (Figure 8). All three between chromosome group were significant (*P* < 0.001). Furthermore, the windows with extremely high diversity and windows with extreme values of Tajima’s *D* statistic were examined for their SNP density.

**Figure 7.**
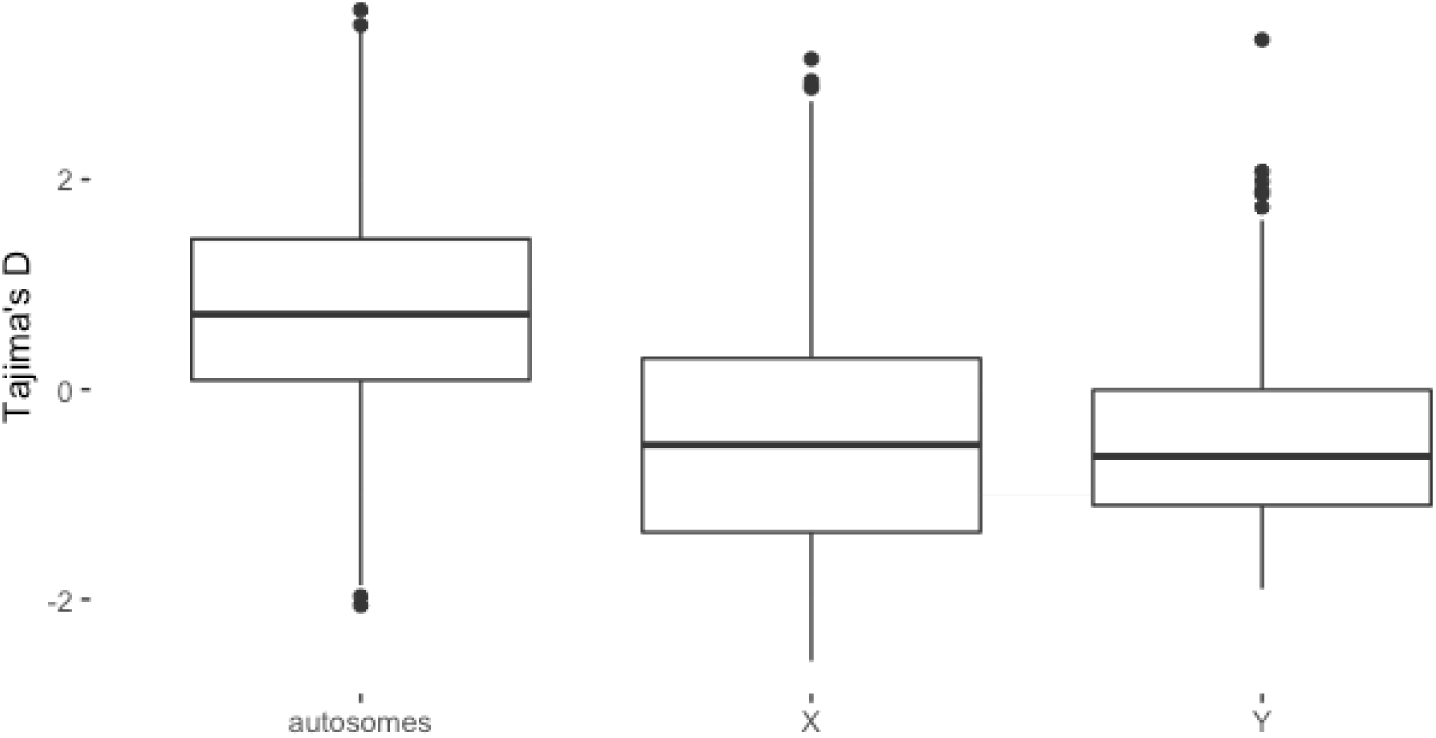
Tajima’s *D* for autosomes, the BTX chromosome, and the BTY chromosome.

**Figure 8.**
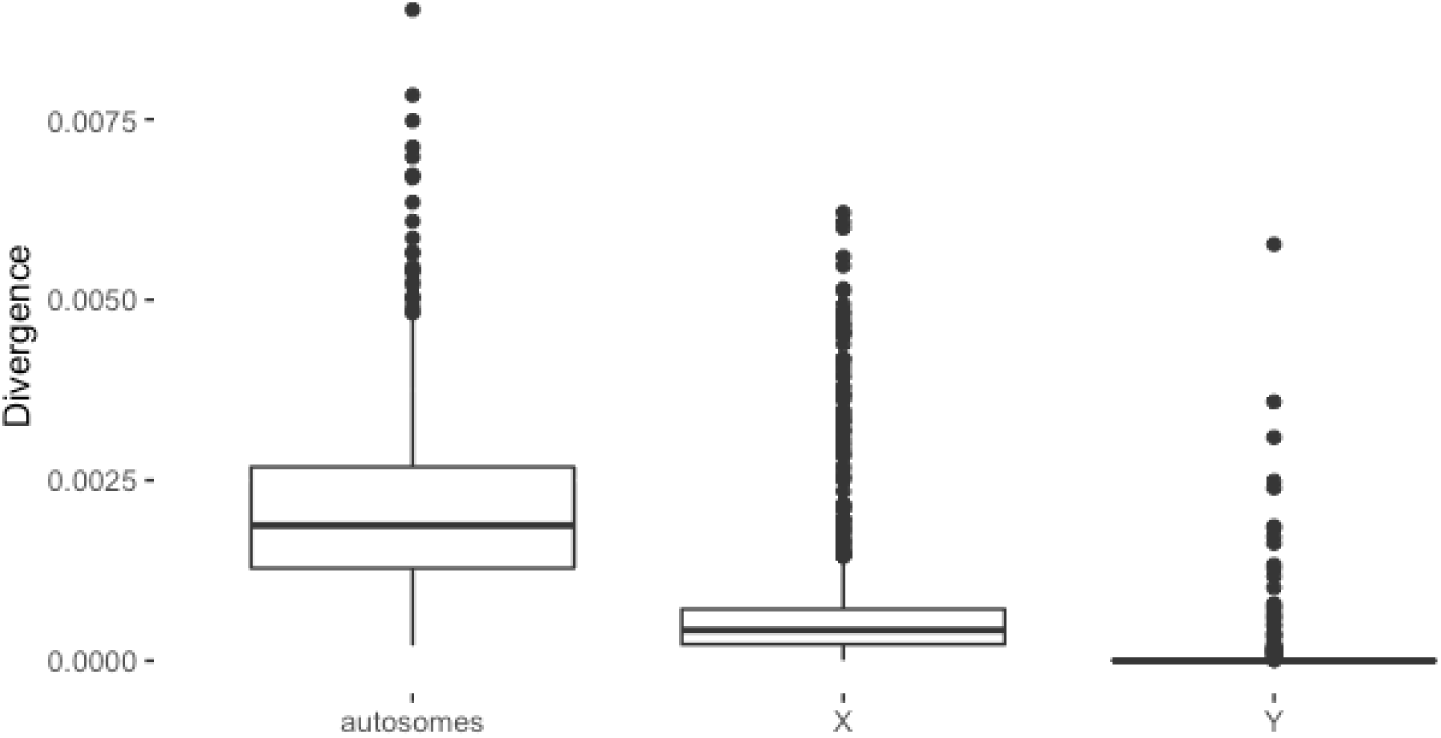
Nucleotide divergence for autosomes, the BTX chromosome, and the BTY chromosome.

## Discussion

Our goal with this study was to compare patterns of genetic variation between autosomes, the BTX and the BTY. A high percent of mapped reads, high percent of properly mapped paired reads and the fact hat the majority of individual genomes was sequenced with a genome average coverage exceeding twenty demonstrated high quality of our data allowing for reliable inferences. Interestingly, individuals with low percent of mapped reads and low percent of properly mapped reads had high genome average coverage. This implies no correlation between genome average coverage and percent of mapped and properly mapped reads.

Altogether, over 23.6 million SNPs and over 3.7 million InDels were identified, among them 24% of SNPs and 94% of InDels were novel. Such proportion between novel SNPs and InDels can be caused by the fact that many more studies have been related to SNPs and thus the SNP data base is more complete ^5^. A larger number of SNPs than InDels is not an uncommon observation. Genic InDels are highly deleterious to gene function as they can completely alter protein amino acid sequence by changing the open reading frame (frameshift mutation). BTY contains the lowest number of variants, which is not only due to its length. For instance, BTA25 is the shortest autosome, but it contains much higher number of variants. Taking into account the length of each chromosome, we observed that only 0.1% of whole length of the BTY contained SNPs, where it is 0.45% for BTX. On the other hand, the shortest chromosome (BTA25) exhibited the highest SNPs density of 2.58%. One of the factors affecting DNA variation is effective population size (*N*_*e*_). Estimated based on autosomes, the effective population size is higher than based on sex chromosomes – ¾ of autosomal *N*_*e*_ for X and ¼ of autosomal *N*_*e*_ for Y ^6^. The lower *N*_*e*_, the more important role of genetic drift, so we expect that the effect of genetic drift is the highest for the Y chromosome, which furthermore implies lower neutral diversity of Y ^7^. This observation contradicts with the theory of male mutation bias that asummes that more mutations accumulate in male germline due to a greater number of male germline cell divisions, what is especially pronounced for BTY which “spends more time” in male germ line than the other chromosomes ^8^. A difference observed in our study is possibly due to gaps in the BTY chromosome assembly of the bovine genome (i.e. a high number of unknown nucleotides), which humper an accurate comparison of variant density between autosomes, BTX and BTY.

However, more recent studies emphasise that the comparison of mutation patterns on a general, chromosome-wide scale, is not valid due to a strong variation in local mutation rates ^9 10^. Also in our study SNP and InDel distribution showed a non-random pattern, characterised by mutation hotspots with very high variant density, albeit only on autosomes. We observed that SNP density is positively correlated with indel density, which is in agreement with results obtained for humans by Hodgkinson *et al.* ^11^. *Estivill et al.* ^12^ showed that regions with high-density of SNPs are correlated with segmental duplications in the human genome. Varela and Amos ^13^ declared that regions with unusually high recombination rates tend to have high density of SNPs, while Aggarwala and Voight ^14^ estimated the effect of DNA sequence k-mers on SNP probability.

Based on annotation of genic regions, it can be shown that there is a tendency that genic variants on BTX have less severe consequences as compared to Y and autosomes. Fewer extreme variants is consistent with purging due to the hemizygous state in males. On autosomes, these would most of the time be hidden in males due to diploidy. Based on Ka/Ks ratio, we also observed that on the BTY there is tendency to accumulate more nonsynonymous than synonymous substitutions (Ka/Ks = 2.00). In contrast, BTX shows a much lower ratio (Ka/Ks = 0.62). Such differences might arise from tendency for a degeneration of the Y chromosome – Haldane’s rule, which indicates faster male evolution and reason of accumulation of nonsynonymous variants, mainly due to the lack of recombination ^15^. In males, BTX is hemizygous, so any, even slightly deleterious mutation has an effect on the phenotype. For mammalian genomes situation was demonstrated in a simulation study by Mackiewicz *et al.*. ^16^

## Methods

### Material

The material comprised whole genome DNA sequences of 217 individuals representing Holstein (69 bulls and 6 cows), Jersey (41 bulls), Jysk (5 bulls), Rød Dansk Malkerace (46 bulls), Rød Dansk Malkerace anno 1970 (15 bulls and 5 cows), Sortbroget Dansk Malkerace anno 1965 (15 bulls and 5 cows) and Danish Shorthorn (5 bulls and 5 cows) breeds available courtesy of the Center for Quantitative Genetics and Genomics at the Aarhus University.

Whole-genome DNA sequences were obtained by the Illumina HiSeq 2000 Next Generation Sequencing platform and similar short-read sequencing platforms. Access to this data was available via the computer cluster of the Center for Quantitative Genetics and Genomics at Aarhus University.

ARS-UCD1.2_Btau5.0.1Y ^1^ reference genome has been used to processing whole-genome sequence data. This genome represents the latest reference genome of *Bos taurus* additionally assembled with Y chromosome (Btau5.0.1) from Baylor College ^17^. In this study two GFF (General Feature Format) files were used for the annotation of identified variants. Btau_5.0.1 and ARS-UCD1.2 GFF were merged to obtain a complete annotation file for each chromosome. Those files were downloaded from the NCBI database (ftp://ftp.ncbi.nlm.nih.gov/genomes/all/GCF/002/263/795/GCF_002263795.1_ARS-UCD1.2/ and ftp://ftp.ncbi.nlm.nih.gov/genomes/all/GCF/000/003/205/GCF_000003205.7_Btau_5.0.1/). The actual assembly file used was downloaded from https://sites.ualberta.ca/~stothard/1000_bull_genomes/ARS-UCD1.2_Btau5.0.1Y.fa.gz.

### Processing the Next-Generation Sequencing Data

The FastQC software was used to summarize and visualize sequence quality ^18^. Sequence quality metrics measure the probability that a given base is called incorrectly. Low quality bases were trimmed from reads using Trimmomatic ^19^ with options SLIDINGWINDOW:3:15, LEADING:3, TRAILING:3 and a minimum read length of 70 bp. Cleaned reads were aligned to the reference genome by using the Burrows-Wheeler Aligner (BWA) using the MEM algorithm ^20^. After alignment, the flagstat tool in SAMtools ^21^ was used to calculate the percent reads correctly mapped to the reference genome. The genome average coverage, representing the number of time that a base in the reference genome was covered by aligned reads ^22^, was calculated for each individual, using the genomecov tool in bedtools ^23^. It was expressed by coverage 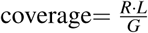, where *R* is the total number of aligned reads, *L* is average read length and *G* is the genome size. The output from BWA-MEM was piped to SAMtools ^21^ and then compressed to the BAM format (Binary Alignment Map) — a binary version of the SAM format. Afterwards, SAMtools fixmate ^21^ was run to adjust the mate-read position. BAM files were sorted using the SAMtools ^21^. PCR duplicates were marked using MarkDuplicates from Picard ^24^. Finally, Base quality scores were recalibrated using Genome Analysis Toolkit ^25^.

### Variant calling

Variant calling allows the identification differences between analyzed sequences and the reference genome, i.e., of polymorphic sites. First, GATK’s HaplotypeCaller ^25^ was used to create files (gVCF files) which summarize information on sites potentially deviating from the reference. Specifically, the tool identifies genomic regions, so called ActiveRegions, which contain significant differences from the reference genome. Those regions are then processed by HaplotypeCaller. Other variations (non ActiveRegions) are skipped in order to accelerate the analysis. Afterwards, ActiveRegions are used to construct haplotypes by building a De Bruijn-like graph ^26^ and calculate haplotype frequencies. Haplotypes were realigned against the reference haplotype using the Smith-Waterman algorithm ^27^ to identify potentially polymorphic sites. Then a matrix of likelihoods of haplotypes given the DNA sequence of reads was calculated using Hidden Markov Models. Thereafter, HaplotypeCaller assigned the most likely genotypes ^25^. Phred-scaled confidence threshold of 30 is typically used in order to control the false-positive variant ratio. The last step of variant calling comprised merging gVCF records. For this purpose, GATK GenotypeGVCFs was used ^25^. This step resulted in raw variant call files (VCF), which contained summary information of each detected variant.

After variant calling, the variants were annotated with predicted biological consequences and functions. For each polymorphic variant the associated Sequence Ontology (SO) ^28^ terms, which categorize genomic functions of the coding sequence, were assigned. For this purpose, the snpEff software was used ^29^. The program uses information included into the annotation file (GFF) to assign an annotation to each detected variant.

### Statistical analysis

#### The Shapiro-Wilk test

The Shapiro-Wilk ^30^ was used to test where a sample was obtained from a normal distribution comparing the following hypotheses:

*H*_0_ The distribution of Tajima’s D and nucleotide divergence follow the normal distribution

*H*_1_ The distribution of Tajima’s D and nucleotide divergence do not follow the normal distribution

The test statistic for Shapiro-Wilk test is as follows:

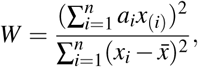

where *a*_*i*_ is the tabulated Shapiro-Wilk coefficient, *x*_(*i*)_ is the *i*th smallest value of the Tajima’s D/nucleotide divergence and 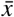 is the mean of Tajima’s D/nucleotide divergence. We reject *H*_0_ at the significance level (*α* = 0.05) if *W* < *W*_*α*_, where *W*_*α*_ is tabulated critical threshold for Shapiro-Wilk.

#### The χ^2^ goodness-of-fit test

The *χ*^2^ goodness-of-fit test was used to check whether the observed variable in the general population follows the uniform distribution. Corresponding hypotheses were defined as follows ^31^:

*H*_0_ The distribution of the observed variable is uniform

*H*_1_ The distribution of observed variable is not uniform

The *χ*^2^ goodness-of-fit statistic is given by ^32^:

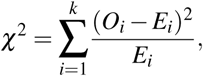

where *O*_*i*_ is the observed count of observations in the *i*^th^ group, *E*_*i*_ is the count of observations expected by the uniform distribution and *k* is the number of groups. Under *H*_0_, the test follows the *χ*^2^ distribution with *k* − 1 degrees of freedom.

#### The Kruskal-Wallis test

The Kruskal-Wallis test is a non-parametric equivalent of the *F* test in the analysis of variance for non-normal data. Corresponding hypotheses are ^33^:

*H*_0_ There is no difference among *k* populations’ median

*H*_1_ At least one population’s median differs from the median of the other populations

The Kruskal-Wallis statistic is given by ^34^:

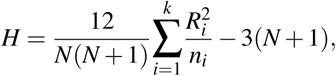

where *N* is the total sample size, *k* is the number of groups, *R*_*i*_ is the sum of ranks in the *i*^th^ group, and *n*_*i*_ is the size of the *i*^th^ group. Under *H*_0_, the test follows the *χ*^2^ distribution with *k* − 1 degrees of freedom where *k* is the number of populations.

#### The Mann-Whitney U test

The Mann-Whitney *U* test is the non-parametric equivalent of the *t*-test for two independent samples with the corresponding hypotheses ^35^:

*H*_0_ There is no difference between two populations’ medians

*H*_1_ There is a difference between two populations’ medians

The statistic is given by ^36; 37^:

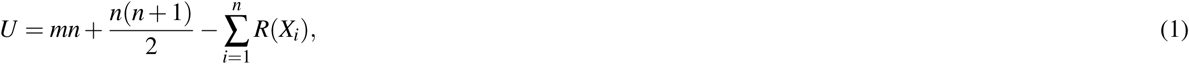

where *n* is the size of first group, *m* is the size of second group, and *R*(*X*_*i*_) is the rank assigned to the first group (Wilcoxon statistic). We reject the null hypothesis *H*_0_ at the significance level *α* if 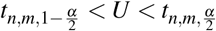, where 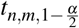 and 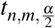 are quantiles of the Mann-Whitney distribution. For large sample sizes, 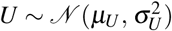, where 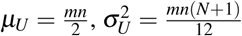, where *N* = *n* + *m*. This test was applied to test multiple, simultaneous hypotheses; therefore the Bonferroni correction was used to account for multiple testing ^38^.

### Genetic statistics

#### Nucleotide divergence

One of the statistic most widely used to measure the degree of polymorphism in chromosome is a nucleotide divergence (*π*). It is a measure proposed by Nei and Li ^39^. *π* quantifies the nucleotide diversity among several sequences. In our case, we estimated *π* along each chromosome in 100 kb non-overlapping windows. The estimator 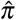 of nucleotide divergence calculated based on VCFs files is as follows:

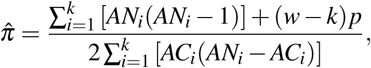

where *k* is the number of variants within a given interval, *AN*_*i*_ is the total number of alleles in called genotypes of *i*^th^ variant, *w* denotes the size of interval (window) [100 kb], *p* is the number of pairwise chromosome comparisons, and *AC*_*i*_ is the allele count in genotypes of *i*^th^ variant (for each alternate allele).

#### Tajima’s D

The second statistic used in this study is Tajima’s *D*. It allows for detection of the evidence of selection. Tajima ^40^ proposed this statistic as a measure of a rate of a random (neutral) evolution of DNA sequence. In this study, Tajima’s *D* was estimated over a 100 kb non-overlapping windows as follows:

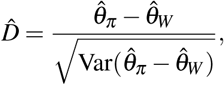

where 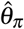 is the average number of pairwise differences given by 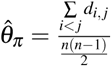. Here, *d*_*i, j*_ represents the number of differences between individual *i* and *j*, 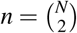 is the number of pairwise sequences comparisons with *N* being the number of individuals. 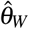 is Watterson’s estimator of the expected number of segregating sites under neutrality 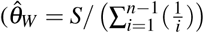, where *S* is the number of sites that segregate in the sample).

*D* > 0 indicates population reduction or balancing selection, *D* = 0 indicates neutral mutations, and *D* < 0 indicates population expansion or purifying and positive selection ^41^. The VCFtools ^42^ was used to calculate the Tajima’s D statistic.

#### K_a_/K_s_ ratio

The *K*_*a*_*/K*_*s*_ ratio was used as a measure of the selection pressure. Based on this ratio we are able to compare natural selection pressure on proteins between chromosomes ^43^. *K*_*a*_ represents the ratio of the number of nonsynonymous substitutions per non-synonymous site, while *K*_*s*_ is the number of synonymous substitutions per synonymous site. *K*_*a*_*/K*_*s*_ equal to one indicates neutral selection, positive ratio is equivalent to positive selection, while negative ratio indicates purifying selection. The output of the SnpEff ^29^ software was used to calculate *K*_*a*_*/K*_*s*_ ratio.

### Computing environment

All programs, and scripts were written in the bash command language were and executed on the Genomics High Performance Cluster (GHPC) at the Center for Quantitative Genetics and Genomics at Aarhus University. The computing unit was the Red Hat Enterprise Linux Client release 4.8.5-28 (Centos) with 250 GiB/node of memory and 6 Intel Core Processor CPUs.

All statistical analyzes were done using R (version 5.1) ^44^ in RStudio ^45^ and visualized using the ggplot2 R package ^46^.

## Data Availability Statement

The subset of the whole-genome data used in this study is available at SRP039339 under PRJNA238491. For the remaining data, the Board of the 1000 Bull Genome Project Consortium should be contacted. Whole-genome sequences from Aarhus University are available only upon agreement with the breeding organization and should be requested directly from the authors.

## Author contributions statement

B.G. and J.S conceived and conducted the experiment, B.C. analyzed the results and wrote the manuscript. All authors reviewed the manuscript.

## References

1. Rosen, B. D. et al. Modernizing the Bovine Reference Genome Assembly. Mol. Genet. 3, 802 (2018).

2. Zimin, A. V. et al. A whole-genome assembly of the domestic cow, bos taurus. Genome Biol. 10, R42, DOI: 10.1186/gb-2009-10-4-r42 (2009).

3. Daetwyler, H. D. et al. Whole-genome sequencing of 234 bulls facilitates mappig of monogenic and complex traits in cattle. Nat. Genet. 46, 858–865 (2014).

4. Chang, T.-C., Yang, Y., Retzel, E. F. & Liu, W.-S. Male-specific region of the bovine y chromosome is gene rich with a high transcriptomic activity in testis development. Proc. Natl. Acad. Sci. 110, 12373–12378, DOI: 10.1073/pnas.1221104110 (2013).

5. Choi, J.-W. et al. Massively parallel sequencing of chikso (korean brindle cattle) to discover genome-wide SNPs and InDels. Mol. Cells 36, 203–211, DOI: 10.1007/s10059-013-2347-0 (2013).

6. VanBuren, R. et al. Extremely low nucleotide diversity in the x-linked region of papaya caused by a strong selective sweep. Genome Biol. 17, DOI: 10.1186/s13059-016-1095-9 (2016).

7. Hellborg, L. Low levels of nucleotide diversity in mammalian y chromosomes. Mol. Biol. Evol. 21, 158–163, DOI: 10.1093/molbev/msh008 (2003).

8. Goetting-Minesky, M. P. & Makova, K. D. Mammalian male mutation bias: Impacts of generation time and regional variation in substitution rates. J. Mol. Evol. 63, 537–544, DOI: 10.1007/s00239-005-0308-8 (2006).

9. Duret, L. Mutation patterns in the human genome: More variable than expected. PLOS Biol. 7, 1–3, DOI: 10.1371/journal.pbio.1000028 (2009).

10. Amos, W. Even small SNP clusters are non-randomly distributed: is this evidence of mutational non-independence? Proc. Royal Soc. B: Biol. Sci. 277, 1443–1449, DOI: 10.1098/rspb.2009.1757 (2010).

11. Hodgkinson, A., Ladoukakis, E. & Eyre-Walker, A. Cryptic variation in the human mutation rate. PLoS Biol. 7, e1000027, DOI: 10.1371/journal.pbio.1000027 (2009).

12. Estivill, X. Chromosomal regions containing high-density and ambiguously mapped putative single nucleotide polymorphisms (SNPs) correlate with segmental duplications in the human genome. Hum. Mol. Genet. 11, 1987–1995, DOI: 10.1093/hmg/11.17.1987 (2002).

13. Varela, M. A. & Amos, W. Heterogeneous distribution of SNPs in the human genome: Microsatellites as predictors of nucleotide diversity and divergence. Genomics 95, 151–159, DOI: 10.1016/j.ygeno.2009.12.003 (2010).

14. Aggarwala, V. & Voight, B. F. An expanded sequence context model broadly explains variability in polymorphism levels across the human genome. Nat. Genet. 48, 349–355, DOI: 10.1038/ng.3511 (2016).

15. Charlesworth, D., Charlesworth, B. & Marais, G. Steps in the evolution of heteromorphic sex chromosomes. Heredity 95, 118–128, DOI: 10.1038/sj.hdy.6800697 (2005).

16. Mackiewicz, D., Posacki, P., Burdukiewicz, M. & Błażej, P. Role of recombination and faithfulness to partner in sex chromosome degeneration. Sci. Reports 8, DOI: 10.1038/s41598-018-27219-1 (2018).

17. Bellott, D. W. et al. Mammalian Y chromosomes retain widely expressed dosage-sensitive regulators. Nature 508, 494–499 (2014).

18. Andrews, S. FastQC A Quality Control tool for High Throughput Sequence Data. http://www.bioinformatics.babraham.ac.uk/projects/fastqc/ (2014).

19. M Bolger, A., Lohse, M. & Usadel, B. Trimmomatic: A Flexible Trimmer for Illumina Sequence Data. Bioinforma. (Oxford, England) 30 (2014).

20. Li, H. & Durbin, R. Fast and accurate short read alignment with burrows–wheeler transform. Bioinformatics 25, 1754–1760, DOI: 10.1093/bioinformatics/btp324 (2009). /oup/backfile/content_public/journal/bioinformatics/25/14/10.1093/bioinformatics/btp324/2/btp324.pdf.

21. Li, H. et al. The Sequence Alignment/Map format and SAMtools. Bioinformatics 25, 2078–9, DOI: 10.1093/bioinformatics/btp352 (2009).

22. Sims, D., Sudbery, I., Ilott, N. E., Heger, A. & Ponting, C. P. Sequencing depth and coverage: key considerations in genomic analyses. Nat. Rev. Genet. 15, 121–132, DOI: 10.1038/nrg3642 (2014).

23. Quinlan, A. R. & Hall, I. M. BEDTools: a flexible suite of utilities for comparing genomic features. Bioinformatics 26, 841–842, DOI: 10.1093/bioinformatics/btq033 (2010).

24. Picard. https://broadinstitute.github.io/picard/.

25. McKenna, A. et al. The Genome Analysis Toolkit: a MapReduce framework for analyzing next-generation DNA sequencing data. Genome Res. 20, 1297–303, DOI: 10.1101/gr.107524.110 (2010).

26. Bruijn, de N. A combinatorial problem. Proc. Sect. Sci. Koninklijke Nederlandse Akademie van Wetenschappen te Amsterdam 49, 758–764 (1946).

27. Smith, T. & Waterman, M. Identification of common molecular subsequences. J. Mol. Biol. 147, 195–197, DOI: https://doi.org/10.1016/0022-2836(81)90087-5 (1981).

28. Eilbeck, K. et al. The sequence ontology: a tool for the unification of genome annotations. Genome Biol. 6, R44, DOI: 10.1186/gb-2005-6-5-r44 (2005).

29. Cingolani, P. et al. A program for annotating and predicting the effects of single nucleotide polymorphisms, snpeff: Snps in the genome of drosophila melanogaster strain w1118; iso-2; iso-3. Fly 6, 80–92 (2012).

30. Shapiro, S. S. & Wilk, M. B. An analysis of variance test for normality (complete samples). Biometrika 52, 591–611, DOI: 10.1093/biomet/52.3-4.591 (1965).

31. Agresti, A. An Introduction to Categorical Data Analysis (John Wiley & Sons, Inc., 2007).

32. Pearson, K. X. on the criterion that a given system of deviations from the probable in the case of a correlated system of variables is such that it can be reasonably supposed to have arisen from random sampling. The London, Edinburgh, Dublin Philos. Mag. J. Sci. 50, 157–175, DOI: 10.1080/14786440009463897 (1900).

33. Vehkalahti, K. Kruskal-wallis test. In The Concise Encyclopedia of Statistics, 288–290, DOI: 10.1007/978-0-387-32833-1_216 (Springer New York, 2008).

34. Kruskal, W. H. & Wallis, W. A. Use of ranks in one-criterion variance analysis. J. Am. Stat. Assoc. 47, 583–621, DOI: 10.1080/01621459.1952.10483441 (1952).

35. Vehkalahti, K. Mann–whitney test. In The Concise Encyclopedia of Statistics, 327–329, DOI: 10.1007/978-0-387-32833-1_243 (Springer New York, 2008).

36. Wilcoxon, F. Individual comparisons by ranking methods. Biom. Bull. 1, 80, DOI: 10.2307/3001968 (1945).

37. Mann, H. B. & Whitney, D. R. On a test of whether one of two random variables is stochastically larger than the other. The Annals Math. Stat. 18, 50–60, DOI: 10.1214/aoms/1177730491 (1947).

38. Dunnett, C. W. A multiple comparison procedure for comparing several treatments with a control. J. Am. Stat. Assoc. 50, 1096–1121, DOI: 10.1080/01621459.1955.10501294 (1955).

39. Nei, M. & Li, W. H. Mathematical model for studying genetic variation in terms of restriction endonucleases. Proc. Natl. Acad. Sci. 76, 5269–5273, DOI: 10.1073/pnas.76.10.5269 (1979).

40. Tajima, F. Statistical method for testing the neutral mutation hypothesis by dna polymorphism. Genetics 123, 585–595 (1989). http://www.genetics.org/content/123/3/585.full.pdf.

41. Ezaz, T. & Edwards, S. V. Editorial: Evolutionary feedbacks between population biology and genome architecture. Front. Genet. 9, DOI: 10.3389/fgene.2018.00329 (2018).

42. Danecek, P. et al. The variant call format and VCFtools. Bioinformatics 27, 2156–2158, DOI: 10.1093/bioinformatics/btr330 (2011).

43. Hurst, L. D. The ka/ks ratio: diagnosing the form of sequence evolution. Trends Genet. 18, 486–487, DOI: 10.1016/s0168-9525(02)02722-1 (2002).

44. R Core Team. R: A Language and Environment for Statistical Computing (2018).

45. RStudio Team. RStudio: Integrated Development Environment for R (2016).

46. Wickham, H. ggplot2: Elegant Graphics for Data Analysis (Springer-Verlag New York, 2016).

